# Heterogeneous transmission estimation and strategy optimization for Chikungunya: a vector-borne modeling study differentiating age and sex

**DOI:** 10.64898/2026.04.13.718188

**Authors:** Jiahui Li, Zeyu Zhao, Jia Rui, Jianguo Zhao, Qixuan Luo, Kangguo Li, Wentao Song, Sandra Perez, Roger Frutos, Yanhua Su, Qiuping Chen, Tianxin Xiang, Tianmu Chen

**Author notes:** Corresponding author. Email: Tianxin Xiang,; Tianmu Chen,; Qiuping Chen,. These authors contributed equally: Jiahui Li; Zeyu Zhao; Jia Rui.

## Abstract

Against the backdrop of global climate change and accelerating population mobility in 2025, chikungunya fever (CHIKF) exhibited a trend of worldwide spread, significantly increasing the difficulty of controlling tropical mosquito-borne diseases. To enhance the precision of intervention strategies, this study developed an age- and sex-structured human–mosquito interaction dynamic model based on data from the largest CHIKF outbreak ever recorded in China, and conducted a targeted analysis of prevention and control strategies. By decomposing the basic reproduction number and examining population heterogeneity, asymptomatic males aged 15–59 years were identified as the core transmission group. Optimal control analysis revealed that the synergistic implementation of three measures— reducing the effective human-to-mosquito transmission rate, reducing the effective mosquito-to-human transmission rate, and suppressing mosquito population density—could reduce the overall infection rate by 95.7586%. Among these, mosquito population suppression should be prioritized as a universal core strategy; however, its protective effect on females aged 60 years and above was relatively weak, warranting particular attention. The study further demonstrated that asymmetric intensity combinations targeting these three intervention pathways—such as intensity profiles of “10%, 90%, 90%” or “60%, 80%, 90%”—could achieve effective outbreak control. This research elucidates population-specific transmission patterns and key pathways for intervention intensity, providing a theoretical and strategic foundation for the precise control of mosquito-borne diseases. It also provides actionable operational insights to support rapid response and strategy optimization for future emerging outbreaks.

**Author summary:** CHIKF is a mosquito-borne viral disease that is gradually spreading from tropical regions to other areas. To achieve more precise control of this disease, we developed an age- and sex-structured analytical model based on the largest CHIKF outbreak in China, aiming to provide a scientific basis for responding to potential future outbreaks with inherent uncertainties. The study found that asymptomatic males aged 15–59 years were the primary drivers of transmission and should be prioritized as a key population for reducing viral spread in prevention efforts. When evaluating the effectiveness of different intervention strategies, females aged 60 years and above were the least affected by the implemented measures, indicating that this group should strengthen personal protection to lower their infection risk. Among all control measures, mosquito suppression was the most effective, suggesting that vector control strategies should be prioritized in future outbreak responses.

## Introduction

CHIKF is an acute infectious disease caused by the chikungunya virus (CHIKV) and transmitted through the bites of *Aedes* mosquitoes, primarily *Aedes aegypti* and *Aedes albopictus* [*1-2*]. The disease has an incubation period typically ranging from 3 to 7 days. The general population is susceptible to CHIKV, and infection is thought to confer long-lasting immunity. Sources of transmission include acute-phase patients, asymptomatic individuals, and virus-carrying mosquitoes, with most patients remaining contagious within the first week of symptom onset (*3*). In 2025, a local clustered outbreak of CHIKF, triggered by imported cases, occurred again in Foshan City, Guangdong Province, representing the largest epidemic event of its kind recorded in China to date. In the early stage of the outbreak, the local government responded promptly by issuing “A Letter to All Citizens” on July 18, calling for public participation in mosquito prevention and control activities. This announcement marked the official transition from routine prevention and control measures to an emergency response state, initiating a comprehensive containment phase.

By the end of 2024, more than 110 countries had reported local or imported cases of CHIKF. In the first seven months of 2025 alone, over 240,000 cases and 90 deaths were recorded globally (*4*). Modeling studies project a long-term global average annual infection risk of 0.012 (95% UI: 0.007-0.019), corresponding to approximately 14.4 million (95% UI: 11.0-17.8 million) annual infections worldwide (*5*), underscoring both the persistent transmission pressure and the urgency of control efforts. As a region with extensive distribution of *Aedes albopictus* and prolonged mosquito activity seasons, China possesses natural conditions conducive to rapid virus spread, facing a long-term potential risk of “imported cases triggering local transmission.” Since the first imported case from Sri Lanka was detected in 2008 (*6*), a total of 61 CHIKF public health emergencies were reported nationwide from 2010 to 2019, 56 of which involved single imported cases (*7*). However, local clustered outbreaks—such as those in Dongguan, Guangdong in 2010 (*8*) and Ruili, Yunnan in 2019 (*9*)—highlight that the risk of indigenous transmission cannot be overlooked.

Mathematical models play a crucial role in elucidating the transmission dynamics of vector-borne diseases and guiding precision control strategies. During the first CHIKF outbreak in Guangdong Province in 2010, simulation studies based on ordinary differential equations demonstrated that comprehensive interventions focusing on mosquito control were significantly more effective than strategies relying solely on human-to-mosquito transmission control (*10*). In the early stage of the 2025 outbreak in Foshan, Guangdong, which was triggered by an imported case and led to local transmission, Zhao et al. (*11*) employed mathematical modeling to estimate the basic reproduction number as 7.28, indicating extremely high transmission potential. Further model-based dissection of the transmission chain revealed that the “human-to-mosquito” pathway contributed substantially more than the “mosquito-to-human” pathway, serving as the primary driver of the outbreak. Among different types of infected individuals, symptomatic cases played a particularly prominent role in transmission. A cohort study on CHIKF outbreaks in Europe indicated that the 45-64 age group faces the highest infection risk, with female cases consistently outnumbering males (*12*). This gender disparity may be linked to differences in social activities, exposure opportunities, and immune responses between sexes.

Age- and sex-structured dynamic models have been widely applied in animal population studies (*13-14*), with most models utilizing partial differential equations to characterize population structure (*15-17*). In the field of vector-borne diseases, such structured models are also progressively developing. Forouzannia et al. (*18*) developed an age-structured deterministic model to evaluate the impact of antimalarial drugs on transmission dynamics. Their results indicated that, in the absence of drug-induced resistance, the model exhibited competitive exclusion—where *Plasmodium* strains with higher basic reproduction numbers suppressed others, driving them toward extinction. Aimée Uwineza et al. (*19*) further established a malaria transmission dynamic model incorporating both age and sex structures, highlighting that to control malaria transmission in Rwanda, it is essential to focus on reducing mosquito biting and infection rates. They also emphasized the active participation of women in government-led interventions and strengthening malaria prevention education for children. Recent modeling research on CHIKV in Brazil by Cortes-Azuero et al. (*20*) also indicates that increasing age and female sex are significant risk factors for symptom occurrence, further underscoring the necessity of incorporating demographic structure into transmission models. However, existing research still shows a deficiency in simultaneously integrating age and sex structures into CHIKF transmission dynamic models. This gap limits the ability to quantify the specific roles of different demographic groups in transmission, thereby constraining the development of targeted control strategies.

To address the gaps in existing research, this study proposes an analytical framework for optimizing prevention and control strategies, building upon prior work. This framework was then validated and subjected to empirical analysis using data from the largest CHIKF outbreak recorded in China. We developed a vector-borne transmission dynamic model that incorporates both age and sex structures, dividing the total population into multiple subgroups with distinct epidemiological characteristics. Based on this model framework, we analytically decomposed the basic reproduction number to quantitatively assess the heterogeneous contributions of different sexes, age groups, and infection statuses (symptomatic/asymptomatic) to disease transmission. Furthermore, we introduced optimal control theory to construct a Hamiltonian system with control variables representing mosquito-to-human transmission control, mosquito population suppression, and human-to-mosquito transmission control. By numerically solving this system, we identified the optimal implementation pathways for each intervention under realistic outbreak conditions and comprehensively evaluated the effectiveness of different intervention combinations in controlling epidemic progression.

## Methods

### Structural Vector-borne Model development

This study developed an age- and sex-structured human-vector transmission dynamic model for CHIKF. The human population is stratified into 10*k* compartments, where k denotes the number of age groups. Specifically, for each age group *j*(*j* = 1, …, *k*), the model includes the following compartments:

- *S*_*mj*_/*S*_*fj*_: Susceptible males / females in age group j;
- *E*_*mj*_/*E*_*fj*_: Exposed males / females in age group j;
- *I*_*mj*_/*I*_*fj*_: Symptomatically infectious males / females in age group j;
- *A*_*mj*_/*A*_*fj*_: Asymptomatically infectious males / females in age group j;
- *R*_*mj*_/*R*_*fj*_: Recovered males / females in age group j.

And, the vector component comprises five compartments:

- *S*_*a*_: Susceptible larval mosquitoes;
- *I*_*a*_: Infected larval mosquitoes;
- *S*_*v*_: Susceptible adult mosquitoes;
- *E*_*v*_: Exposed adult mosquitoes;
- *I*_*v*_: Infectious adult mosquitoes.

The model is based on the following assumptions:

I. CHIKV transmission occurs exclusively through cross-species interactions, specifically between vectors and humans (vector-to-human or human-to-vector). Direct human-to-human or vector-to-vector transmission is not considered.
II. The effective transmission rate varies across age and sex groups. The rate for the first male age group (*m*1), denoted as *β*_*vp*_, serves as the baseline. The effective transmission rates for all other groups are derived by multiplying this baseline by the corresponding Incidence Rate Ratio (IRR) (S1 Table).
III. Upon infection, susceptible individuals enter a latent period during which they are not infectious. Subsequently, with probability *q*, they progress after an average latency period of 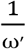, to become asymptomatic infections, eventually recovering at a rate *γ*′. Alternatively, with probability (1 − *q*), they undergo a latency period of 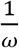 to become symptomatically infectious, then recover at rate *γ*.
IV. A proportion (*n*) of mosquitoes acquire the virus through vertical transmission, influenced by the per capita birth rate of mosquitoes (*a*). This process is modulated by a seasonal effect factor(*c*), making the effective vertical transmission rate *a* · *c* · *n*. Infected larval mosquitoes then develop into infectious adult mosquitoes after an average emergence period of 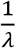, gaining the ability to transmit the virus.
V. Newly reproduced larval mosquitoes, except those vertically infected, enter the susceptible larval mosquito compartment. They develop into susceptible adult mosquitoes after an average emergence period of 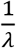. Susceptible adult mosquitoes acquire the virus by biting infectious humans, entering the exposed class. After an average extrinsic incubation period of 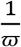, they transition to the infectious class, capable of transmitting the virus.

The model defines the total human population as

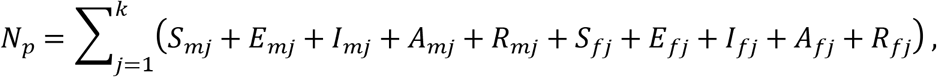

and the total mosquito population as:

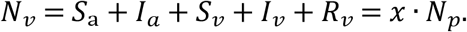

For simplicity in subsequent computations, *N*_*p*_is treated as constant based on its demographic interpretation. Furthermore, the seasonal effect function is defined as:

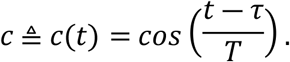

The model structure is illustrated in Fig 1, and the corresponding system of equations is given as follows:

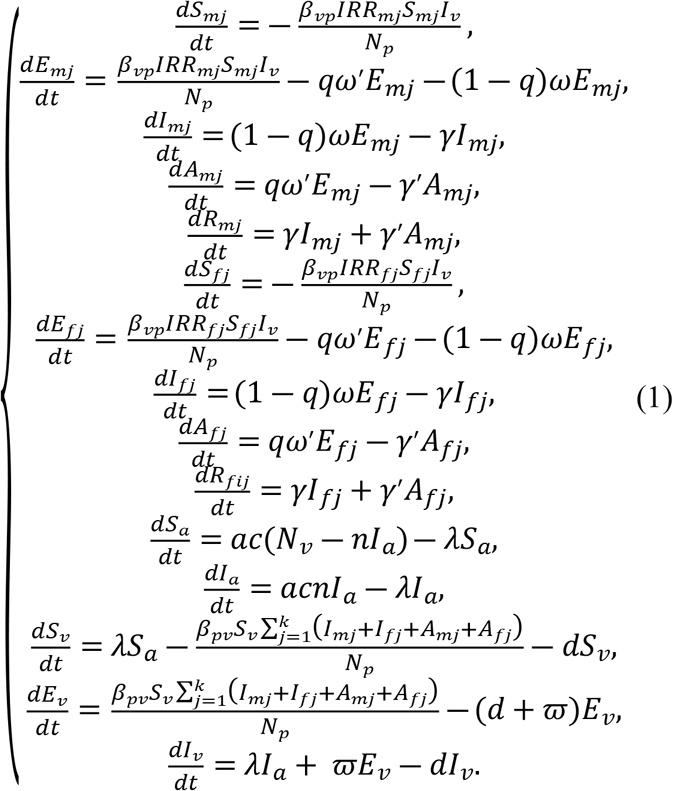

**Fig 1.**
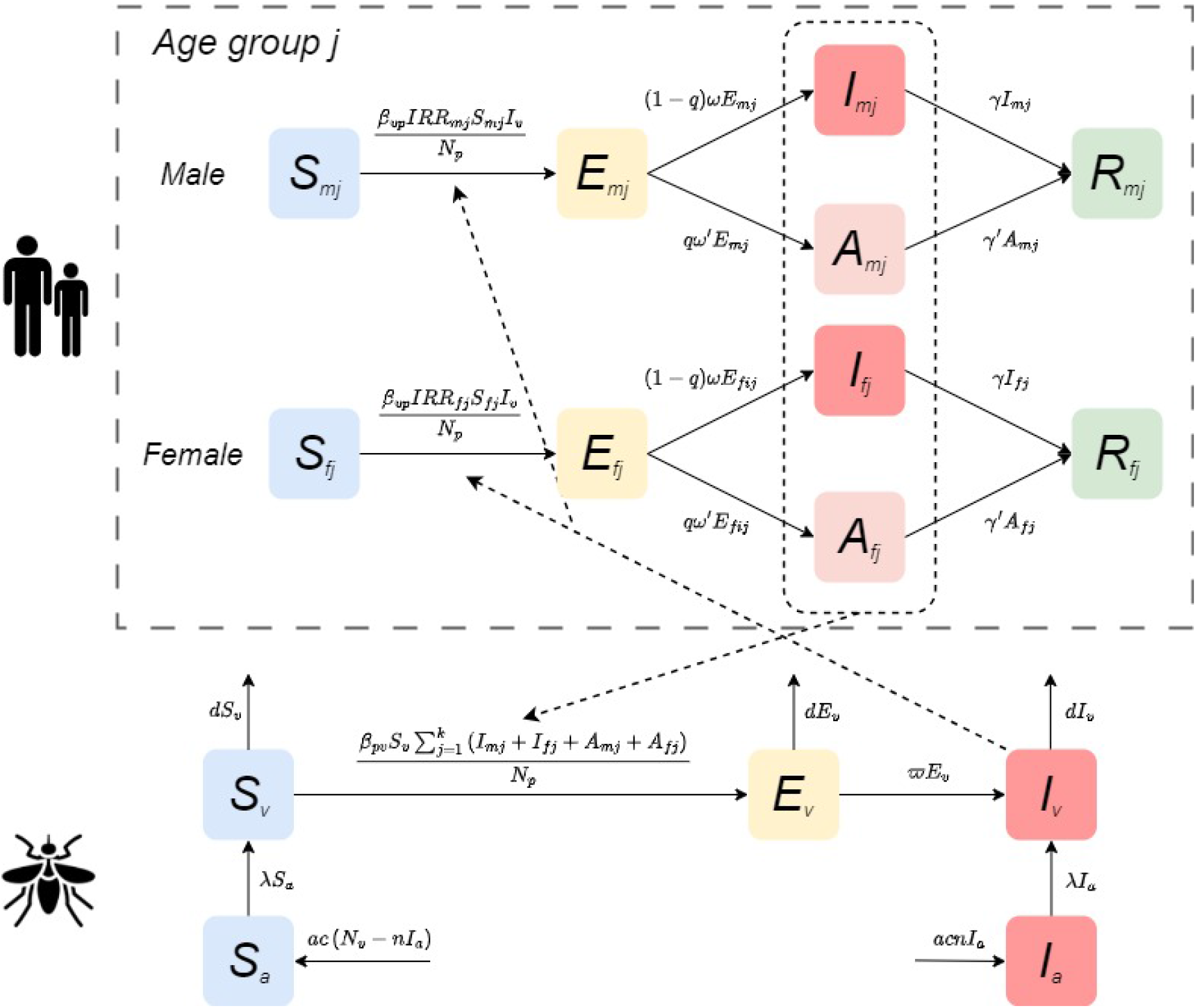
Compartmental structure of the CHIKV transmission model.

### Data and Parameter Collection

The data for this study were sourced from the 2025 CHIKF outbreak records released by the World Health Organization (WHO) (*21*) and the publicly accessible GitHub repository detailing the Foshan outbreak (*22*). In 2025, chikungunya fever presented a globally widespread distribution. According to WHO reports, suspected cases far outnumbered confirmed cases worldwide, suggesting the existence of a substantial number of undiagnosed or asymptomatic infections (Fig 2 b–c). Regionally, the Americas reported the highest number of cases and deaths, accounting for more than half of the global total, while Europe experienced an unusual surge linked to outbreaks in French overseas territories (Fig 2 a, d).

**Fig 2.**
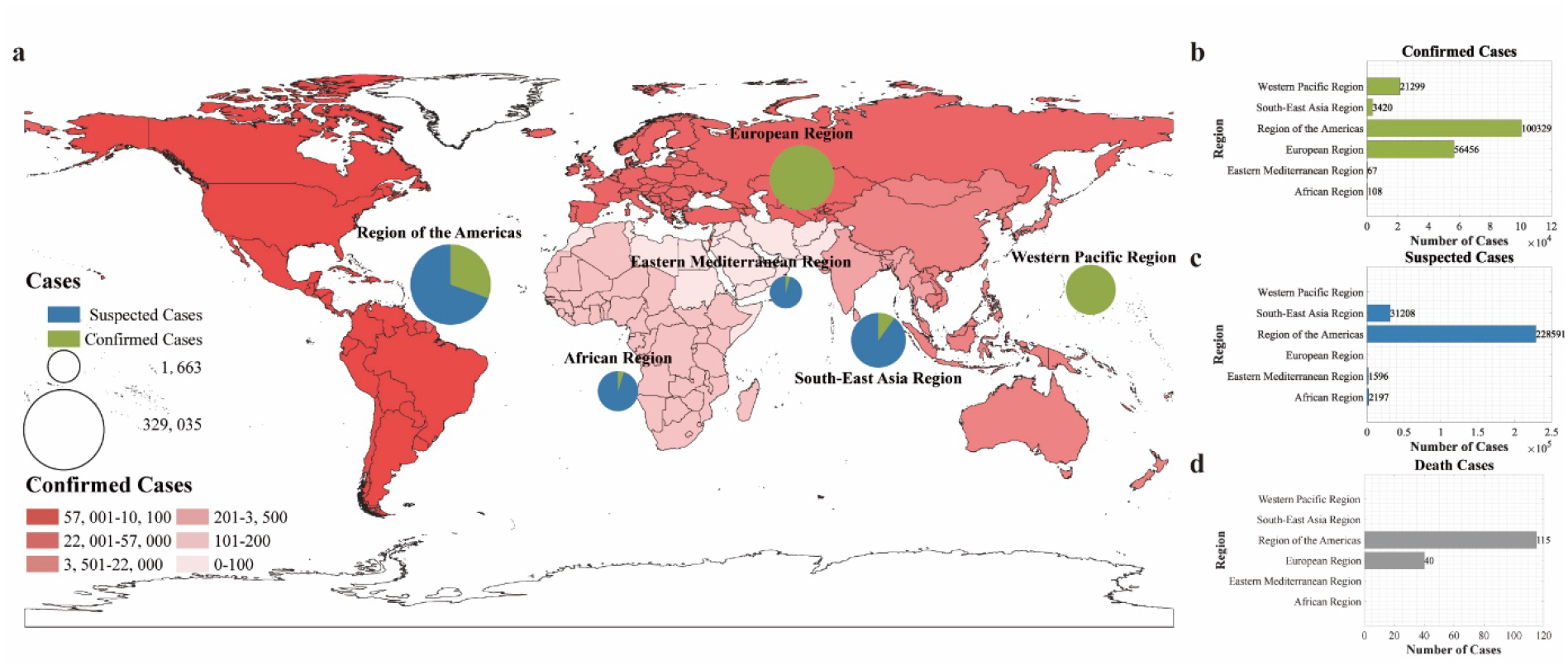
Geographic distribution of suspected and confirmed CHIKF cases and deaths reported to WHO or shared publicly by national ministries of health by region, as of September 2025. **(a)** Burden distribution map by region: color intensity represents case severity, while pie charts show the proportion of suspected versus confirmed cases within each region. **(b)-(d)** Statistics of confirmed cases, suspected cases, and deaths by region, respectively.

To conduct an in-depth analysis of transmission heterogeneity and intervention effectiveness across different population groups, this study selected Foshan City as the study site, which experienced the largest local outbreak in Asia in 2025. Foshan is a typical area with active *Aedes* mosquito populations, and its vector ecology and climatic conditions are representative of southern China and even Southeast Asia. Moreover, the city’s outbreak data are systematic and complete, effectively reflecting human infection patterns in a mosquito-borne transmission context.

During data preprocessing, the epidemic time series was constructed based on the date of symptom onset. According to the available structured data, all cases were stratified by sex (male, female) and age group (children and adolescents: 0-14 years; adults: 15-59 years; older adults: ≥60 years). According to the key phases of the outbreak, the study period was divided into two main intervals for model fitting: a preparedness phase (16 June-18 July 2025) and a containment phase (19 July-18 September 2025). To account for transmission characteristics, the first 7 days of the containment phase (19-25 July) were further defined as an incubation lag phase, allowing assessment of the potential impact of the latent period on outbreak control.

Demographic parameters for the model were derived from population statistics published by the Foshan Municipal Bureau of Statistics (*23*). The parameters IRR were calculated based on the actual number of infections during the model fitting period. Key transmission parameters, including *β*_*vp*_, *β*_*pv*_, and *x*, were estimated using the Particle Markov Chain Monte Carlo (PMCMC) algorithm (S2 Table). The remaining parameter values were obtained from published literature. Detailed descriptions, values, and sources of all model parameters are provided in Table 1.

**Table 1.** Definition and value of parameters in the transmission dynamics model.

| Parameters | Definition | Unit | Value | Range | Source |
| --- | --- | --- | --- | --- | --- |
| $\beta_{vp}$ | Effective transmission rate from mosquitoes to humans | 1 | - | 0-1 | Fitting |
| $\beta_{pv}$ | Effective transmission rate from humans to mosquito | 1 | - | 0-1 | Fitting |
| $x$ | The ratio of mosquitoes to humans total number | 1 | - | 5-15 | Fitting |
| $a$ | Per capita birth rate of mosquitoes | Day<sup>-1</sup> | 0.145 | 0.02-0.27 | Ref. (24-25) |
| $\tau$ | Simulation delay in the initial time in the whole season day | Day | 30 | - | Estimation |
| $T$ | Duration of the daily cycle | Day | 365 | - | Estimation |
| $n$ | The minimum infection rate for vertical transmission | 1 | 0.0181 | 0.0076-0.0286 | Ref. (26-27) |
| $\lambda$ | Rate of mosquito emergence | Day<sup>-1</sup> | 0.0902 | 0.0691-0.1296 | Ref. (28) |
| $d$ | Natural mortality rate of mosquitoes | Day<sup>-1</sup> | 1/7.4 | 1/9.8-1/4.5 | Ref. (29) |
| $\varpi$ | Transition rate of infection in mosquitoes from exposed to infectious | Day<sup>-1</sup> | 1/5.5 | 1/8.2-1/3 | Ref. (30-33) |
| $q$ | Proportion of exposed individuals progressing to asymptomatic individuals | 1 | 0.155 | 0.03-0.28 | Ref. (34-35) |
| $\omega$ | Transition rate of exposed individuals to the symptomatic individuals | Day<sup>-1</sup> | 1/4.5 | 1/7-0.5 | Ref. (34-35) |
| $\omega'$ | Transition rate of exposed individuals to the asymptomatic individuals | Day<sup>-1</sup> | 1/4.5 | 1/7-0.5 | Ref. (34-35) |
| $\gamma$ | Recovery rate of symptomatic individuals | Day<sup>-1</sup> | 1/7 | - | Ref. (36) |
| $\gamma'$ | Recovery rate of asymptomatic individuals | Day<sup>-1</sup> | 1/7 | - | Ref. (36) |

### Parameter Estimation

This study employs the PMCMC method to perform Bayesian inference for the unknown parameter vector *θ* in the dynamic model. The PMCMC framework integrates Particle Filtering (PF) with Markov Chain Monte Carlo (MCMC) sampling, making it particularly suitable for parameter estimation in state-space models characterized by partial observational noise and complex nonlinear dynamics. This approach effectively addresses the challenge of intractable likelihood evaluation inherent in such models (*37*).

The methodological foundation lies in reformulating parameter estimation as the exploration of the posterior distribution *p*(*θ*│*y*_1:*T*_), where *y*_1:*T*_ represents the time-series observational data. According to Bayes’ theorem, the posterior distribution is proportional to the product of the likelihood function *p*(*y*_1:*T*_│*θ*) and the prior distribution *p*(*θ*):

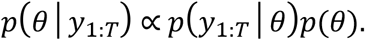

Within this framework, the goodness-of-fit of parameters is defined by their probability density under the posterior distribution. Regions of higher probability density correspond to parameter values that are more consistent with prior knowledge and demonstrate better compatibility with the observed data.

The posterior distribution of parameters is sampled using a constructive Metropolis-Hastings (M-H) algorithm. This method explores the parameter space sequentially by constructing a Markov chain whose stationary distribution is the target posterior distribution (*38*). At each iteration *k*, a candidate parameter value *θ*\* is first generated from a proposal distribution *q*(*θ*│*θ*^(*k*−1)^), typically configured as a Gaussian random walk. To evaluate this candidate, a particle filter is invoked to compute its corresponding marginal likelihood estimate 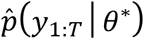. The particle filter sequentially approximates the filtering distribution of the latent states by maintaining a set of weighted particles 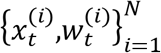, ultimately providing an unbiased estimate of the marginal likelihood:

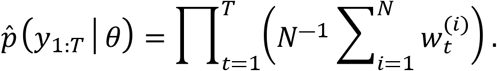

This estimate is embedded within the MCMC sampler, linking parameters to the observed data and driving the exploration of the parameter space.

Based on the output of the particle filter, the algorithm performs an accept-reject step to update the parameters. The acceptance probability *α* for the candidate point *θ*\* is determined by:

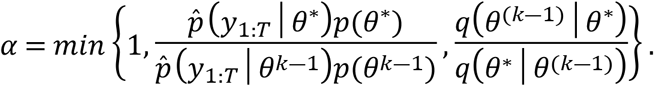

This mechanism ensures that the Markov chain visits regions of higher posterior probability with frequency proportional to their probability density. After a predefined burn-in period, the sequence of samples {*θ*^(*k*)^} generated by the Markov chain can be regarded as an approximation of independent and identically distributed samples from the target posterior distribution. Finally, based on this sample set, statistical inference and uncertainty quantification of the parameter values are completed by calculating the posterior mean E [*θ*|*y*_1:*T*_] or posterior mode as a point estimate, and using quantiles to construct credible intervals (*39*).

### Optimal Control

This study first analyzed the key factors influencing epidemic transmission based on the global regional disease burden data up to September 2025. Building on this, and focusing on human-controllable interventions, we developed an intervention transmission dynamics model that incorporates both human population and vector structural features. The model targets three critical pathways: blocking mosquito-to-human transmission, controlling human-to-mosquito transmission, and suppressing the mosquito population. Guided by Pontryagin’s Maximum Principle (*40*), we formulated an optimal control theoretical model, establishing an analytical framework for optimizing disease control strategies (Fig 3).

**Fig 3.**
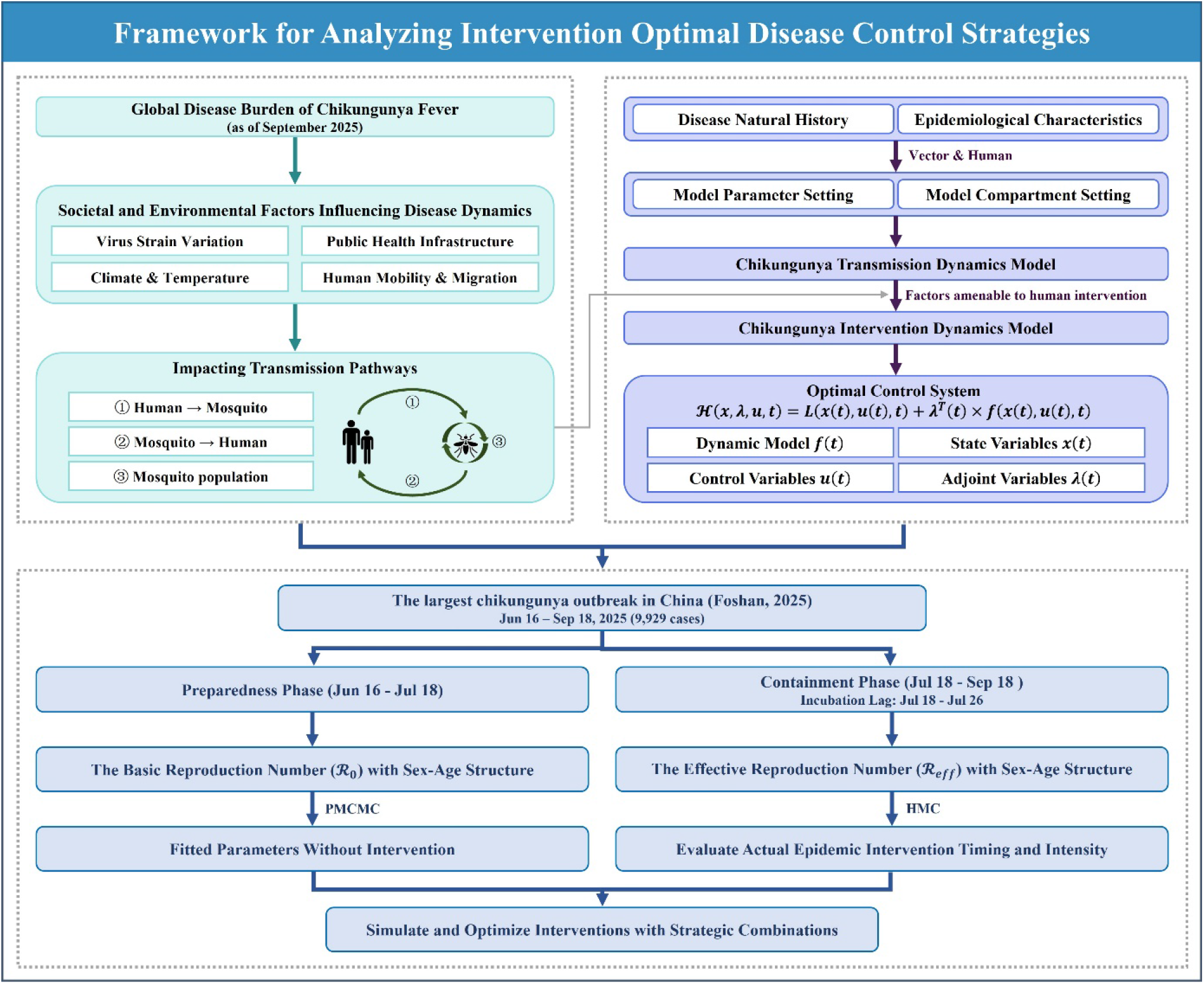
Framework for analyzing intervention optimal disease control strategies.

To validate this framework, we selected the largest CHIKF outbreak recorded in China— the 2025 Foshan epidemic—as a case study for empirical analysis. The Hamiltonian Monte Carlo (HMC) method was employed to derive both analytical and numerical solutions for control strategies within the human-mosquito transmission system, enabling a systematic evaluation of the effectiveness of different intervention combinations. The theoretical core of this method lies in constructing a Hamiltonian function, which transforms the original optimal control problem into solving a coupled system of state and costate equations. This process identifies the optimal control variables that minimize the predefined objective function (*41-42*).

The Hamiltonian function integrates the state equations, costate variables (Lagrange multipliers), and the integrand of the objective function as follows:

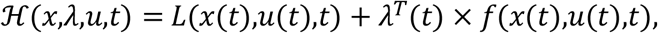

where *x*(*t*) represents the vector of state variables, *u*(*t*) denotes the control variables to be determined, *L*(⋅) corresponds to the integrand (running cost) of the objective function, *f* (⋅) describes the system dynamics, and *λ*(*t*) refers to the costate variables (also known as adjoint variables), with *t* = 0,…,*T*. According to Pontryagin’s Maximum Principle, the optimal control *u*\*(*t*) must minimize (or maximize) the Hamiltonian ℋ over the entire time horizon.

By taking the partial derivative of ℋ with respect to the control variable *u* and setting it to zero, i.e.,

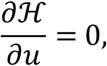

an explicit expression for the optimal control exists and can be derived as a function of the state variables *x*(*t*) and costate variables *λ*(*t*), yielding *u*\* = *u*\*(*x,λ*) (*43-44*).

Solving the aforementioned optimal control system requires simultaneously satisfying both the state equations and the costate equations. The state equation

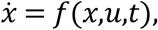

is integrated forward in time with the initial condition *x*(0) = *x*_0_, while the costate equation

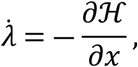

is integrated backward with the terminal condition *λ*(*T*) = 0. To numerically solve this coupled system, we employ the forward-backward sweep method, which iteratively performs the following steps: First, given the current estimate of the control variables, the state equations are integrated forward to obtain the state trajectory; Second, using the current state trajectory, the costate equations are integrated backward to determine the costate trajectory; Finally, the control variables are updated according to the Hamiltonian minimization condition *u*\* = *u*\*(*x,λ*).

To ensure algorithm stability and convergence, a relaxation update strategy is introduced:

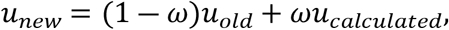

where *ω* ∈ (0,1) is the relaxation (or damping) factor. The updated control variables are then projected onto the admissible control set *U* (e.g., *u*_min_ ≤ *u*(*t*) ≤ *u*_max_), which essentially implements a form of the projected gradient method. The iteration continues until the relative change in the control variables falls below a preset tolerance or the maximum number of iterations is reached.

## Results

### Epidemiological Characteristics

Since the first imported case of CHIKF occurred in Foshan City on June 16, 2025, a total of 9,929 confirmed cases had been recorded by September 18. Among these, 5,171 (52.08%) were male and 4,758 (47.92%) were female, indicating a slightly higher proportion of cases among males (Fig 4d-e). In terms of age distribution, the 15-59 age group was the most affected across both sexes, representing the highest number of infections in each gender subgroup.

**Fig 4.**
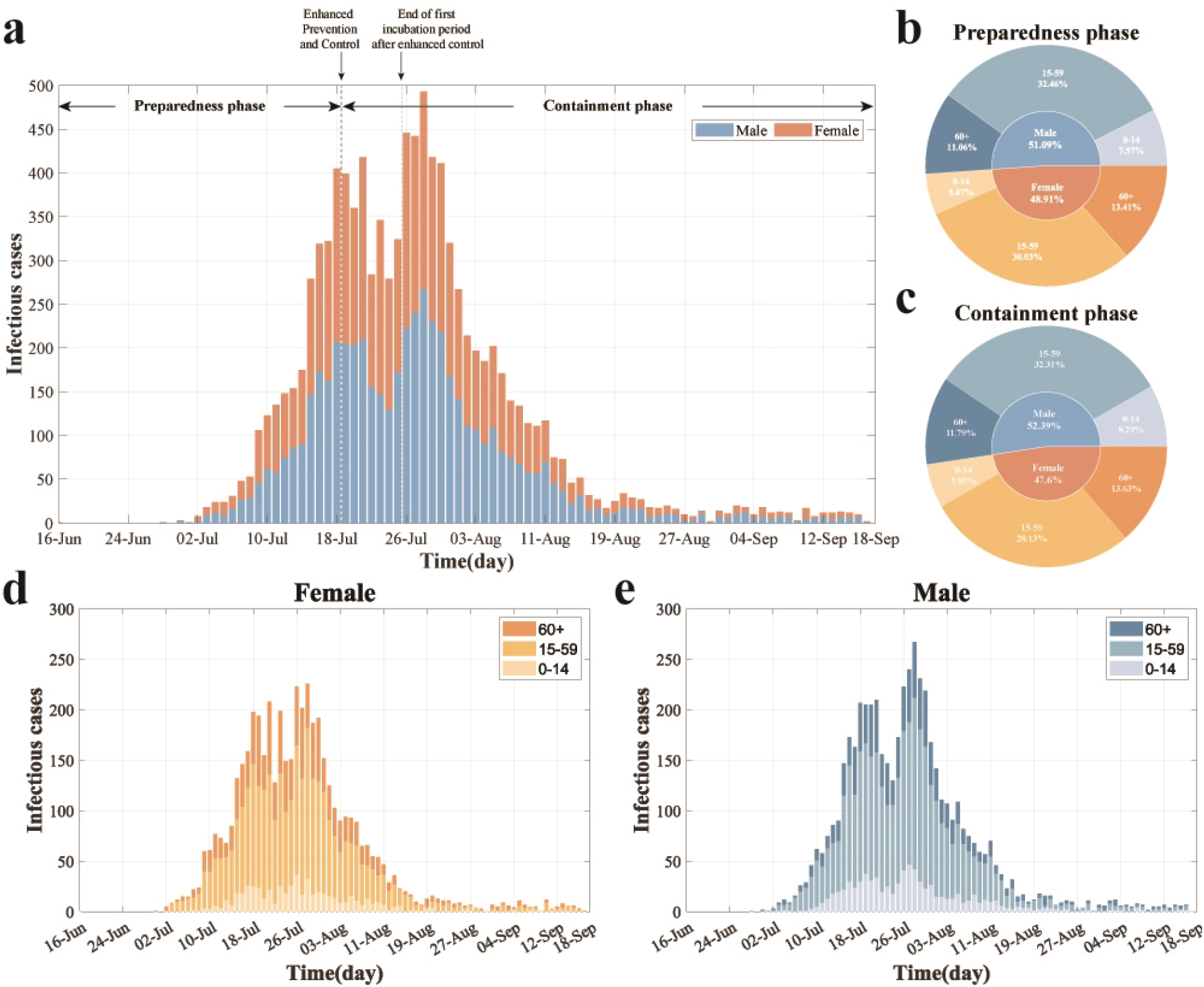
Temporal dynamics and demographic characteristics of the largest recorded CHIKF outbreak in China. **(a)** Sex-stratified daily cumulative reported cases by epidemic phase, with phase divisions annotated in black text and arrows. **(b-c)** Age and sex distribution of incident cases during the preparedness phase (b) and containment phase (c). **(d-e)** Daily cases by age group among females (d) and males (e). Colors represent sex (blue: male; orange: female) and age groups (light to dark shades: 0-14, 15-59, and 60+ years, respectively).

Driven by the implementation of control policies, the epidemic progression exhibited distinct phases and clear intervention-response characteristics (Fig 4a). The first minor peak in daily incidence occurred on July 18, with 405 new cases reported. Following the implementation of containment measures, case numbers were temporarily suppressed. However, after a plateau period reflecting the incubation lag, infections resurged, reaching a peak of 493 new cases on July 28—the highest daily count recorded during the outbreak. Subsequently, the number of new cases declined consistently, indicating that control measures had gradually taken effect and the outbreak had entered a phase of effective containment.

Stratified analysis by phase revealed that during the preparedness phase (June 16-July 18), males accounted for 51.09% of cases. Among all cases in this phase, the 15-59 age group constituted 32.46%, making it the predominant demographic, while individuals aged 60 and above accounted for 11.06%. Females represented 48.91% of cases in this phase, with the 15-59 age group comprising 30.03% of all cases. After transitioning to the containment phase (July 19-September 18), the proportion of male cases slightly increased to 52.39%, with the 15-59 age group remaining the most affected, constituting 32.31% of all cases during this period. This pattern highlights the consistently higher exposure risk and disease burden shouldered by the 15-59 age group throughout the entire epidemic (Fig 4b-c).

### Basic Reproduction Number with Sex-Age Structure

The disease-free equilibrium (DFE) represented by

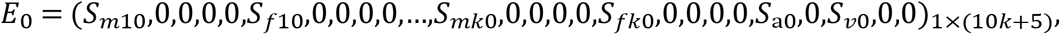

in which 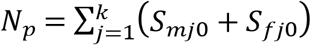 and *N*_*v*_ = *S*_*a*0_ + *S*_*v*0_. We calculated the basic reproduction number ℛ_0_ using the next-generation matrix approach described in (30). The nonlinear terms with new infection ℱ and the outflow term 𝒱 are given by

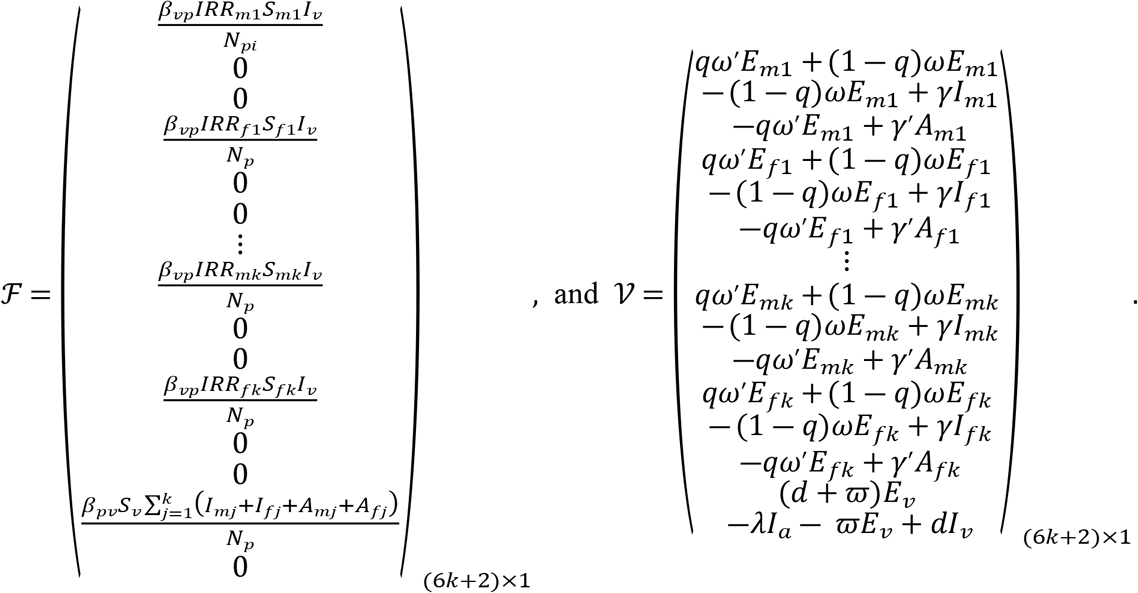

The Jacobian matrices of ℱ and 𝒱 at the DFE *E*_0_ are

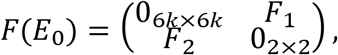

and

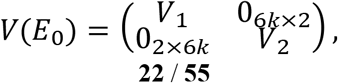

respectively, of which

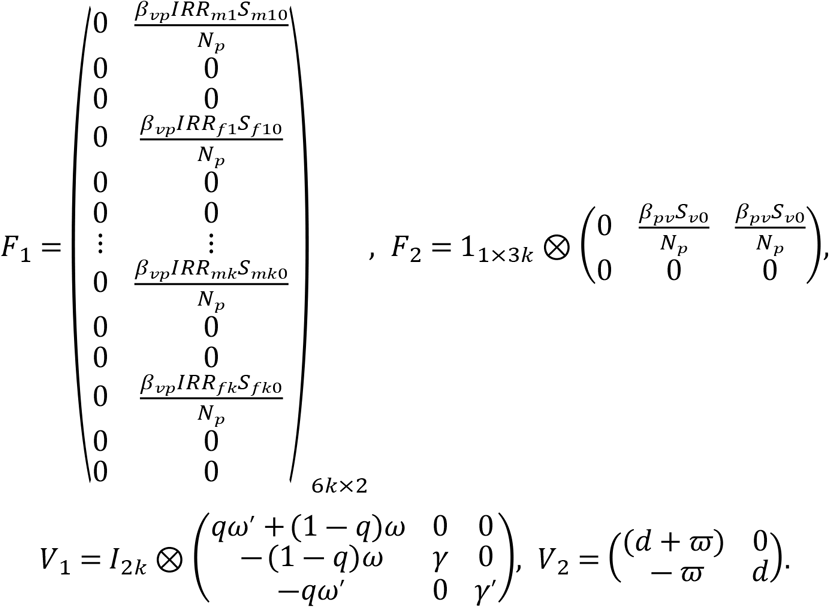

Then, the basic reproduction number ℛ_0_ of model (1) is the spectral radius of the next generation matrix *FV*^−1^(*E*_0_). Direct calculation gives

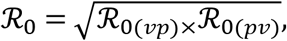

in which,

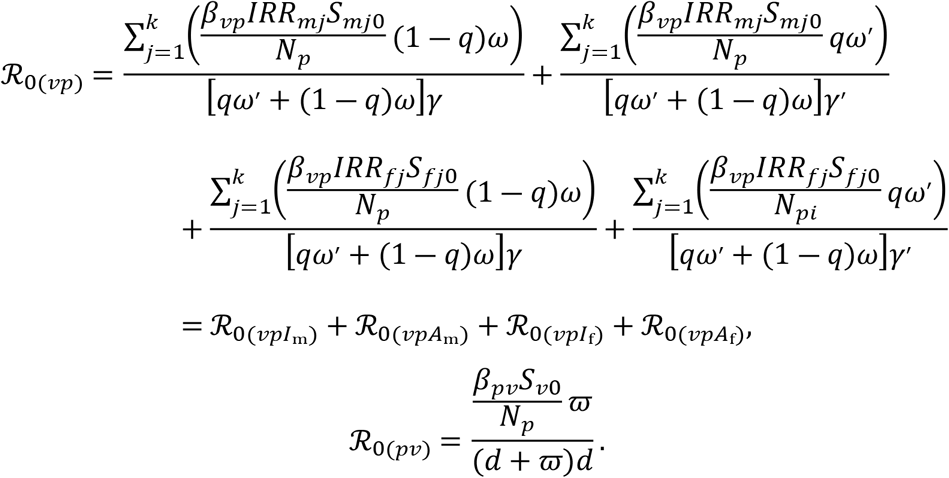

ℛ_0(*pv*)_ represents the average number of secondary human infections produced by a single infected mosquito per unit time in a fully susceptible human population. Specifically, this value can be disaggregated into the sum of four components: the number of secondary symptomatic male infections 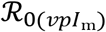, asymptomatic male infections 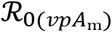, symptomatic female infections 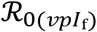, and asymptomatic female infections 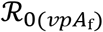. Each of these components is further stratified by j(*j* = 1,…,*k*) age groups. Similarly, 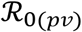 represents the average number of secondary infected mosquitoes produced by a susceptible mosquito, per unit time, through biting the infected population.

In the decomposition analysis of the basic reproduction number, ℛ_0_ was disaggregated into two transmission pathways: “human-to-mosquito” and “mosquito-to-human,” with their geometric mean representing the overall ℛ_0_ of the model (specific values are provided in Table 2). For demographic analysis, the population was divided into three age groups (*k* = 3). The “mosquito-to-human” component was further decomposed into 12 subcomponents, with detailed contributions illustrated in Fig 5.

**Table 2.** Contribution of cross-species transmission pathways to the basic reproduction number (ℛ_0_) in the largest CHIKF outbreak in China.

| Time | $\mathcal{R}_0$ | $\mathcal{R}_{0(vp)}$ | $\mathcal{R}_{0(pv)}$ |
| --- | --- | --- | --- |
| 11 Jun-18 Jul | 10.1235 | 3.6197 | 28.3132 |
| 19 Jul-18 Sep | 1.3865 | 0.8665 | 2.2185 |
| 19 Jul-25 Jul | 1.2934 | 1.0341 | 1.6178 |
| 26 Jul-18 Sep | 1.3243 | 0.8122 | 2.1593 |

**Fig 5.**
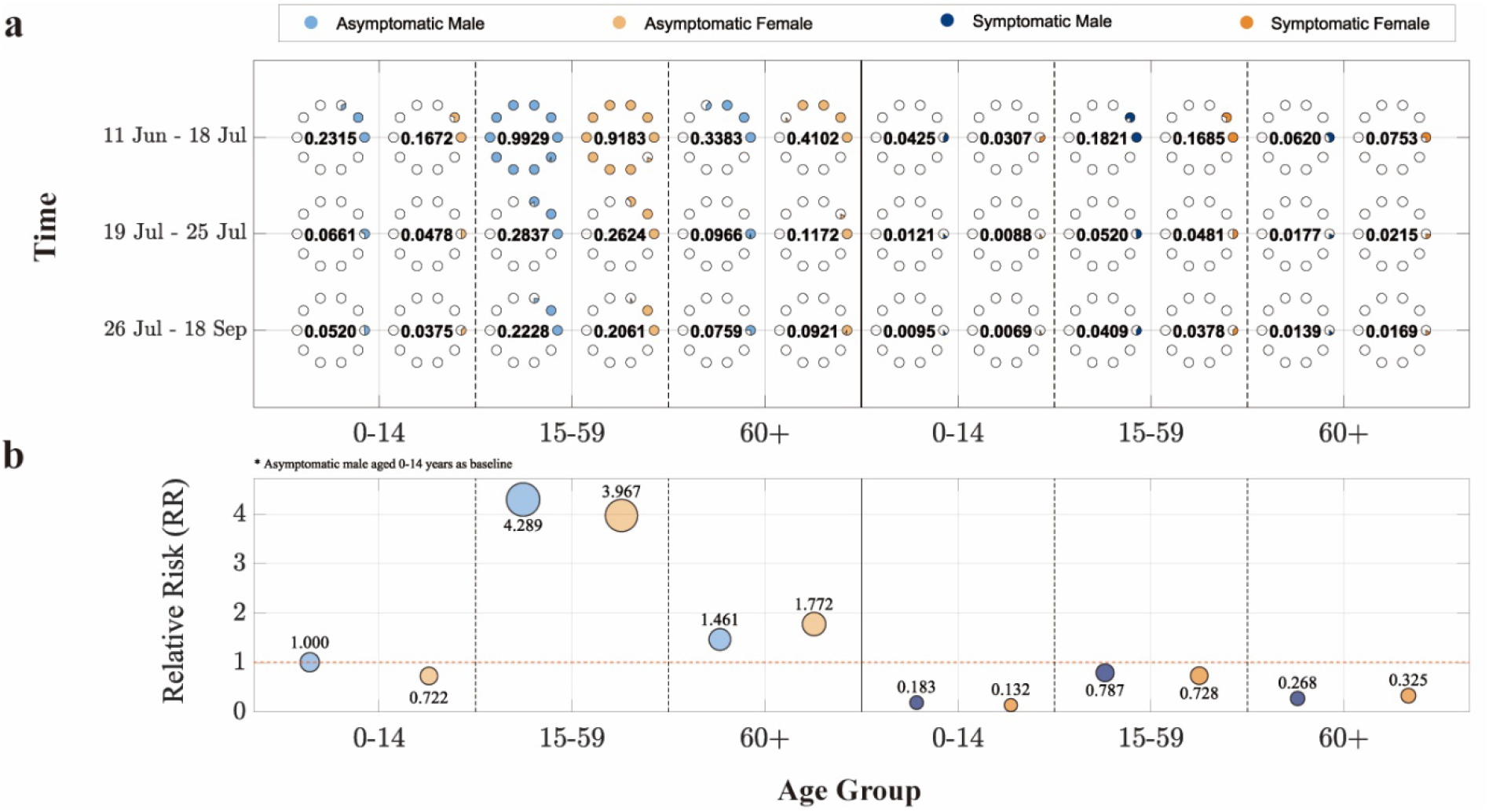
Demographic decomposition and relative risk analysis of the basic reproduction number (ℛ_0_) in the largest CHIKF outbreak in China. **(a)** ℛ_0_ stratified by sex, age group, and infection status (asymptomatic/symptomatic) across different containment phases. The horizontal axis represents age groups, the vertical axis indicates time periods, and point size corresponds to ℛ_0_ values (each hollow circle represents 0.1). **(b)** Relative risk of other population subgroups compared to the reference group (asymptomatic males aged 0-14 years). The solid red line marks the baseline. The vertical position and area of each circle denote the magnitude of relative risk. Blue and orange colors denote male and female cases, respectively, while light and dark blue shades represent asymptomatic and symptomatic infections.

During the preparedness phase, the overall ℛ_0_ remained at a relatively high level. The contribution of “human-to-mosquito” transmission was significantly greater than that of “mosquito-to-human” transmission. The population-wide ℛ_0_ during this phase was 3.6197, exceeding 1, indicating a trend of epidemic expansion. In terms of infection status, asymptomatic individuals contributed more to transmission than symptomatic cases. Among age groups, the 15-59 years group played the most prominent role in transmission. The ℛ_0_ contribution from asymptomatic individuals in this age group reached 0.9929 for males and 0.9183 for females. This was followed by individuals aged 60 and above, among whom the ℛ_0_ contribution from asymptomatic infections was higher in females (0.4102) than in males (0.3383).

During the containment phase, although “human-to-mosquito” transmission remained stronger than “mosquito-to-human” transmission, the overall population ℛ_0_ decreased to 0.8665, falling below 1. This suggests that control measures were effective and the outbreak was brought under improved control. In this phase, asymptomatic individuals in the 15-59 years age group remained the primary contributors to ℛ_0_, with slightly higher values in males (0.2377) than in females (0.2199). When the containment phase was further divided by the incubation lag, the overall model ℛ_0_ during the lag period was slightly lower than in the subsequent period. Further decomposition revealed that although “human-to-mosquito” transmission was lower during the lag period than later, both “mosquito-to-human” transmission and the ℛ_0_ contributions across population subgroups were higher during the lag period. The analysis also reaffirmed that asymptomatic individuals aged 15-59 consistently represented the group with the highest ℛ_0_ contribution throughout.

### HMC-Optimized Strategy Validation

To evaluate the effectiveness of intervention strategies for CHIKF, we referred to the *Global Vector Control Response 2017-2030* issued by the World Health Organization and the *Technical Guidelines for Chikungunya Prevention and Control (2025 Edition*) released by the Chinese Center for Disease Control and Prevention. The various control measures (S5 Table) were categorized into three model-adjustable intervention pathways: controlling mosquito-to-human transmission, controlling human-to-mosquito transmission, and suppressing the mosquito population. These were incorporated into Model (1) as control functions *u*(*t*) = (*u*_1_(*t*),*u*_2_(*t*),*u*_3_(*t*)) (Table 3). A Hamiltonian system was formulated to quantitatively assess the optimal initiation timing and implementation intensity of each intervention in controlling the outbreak.

**Table 3.**
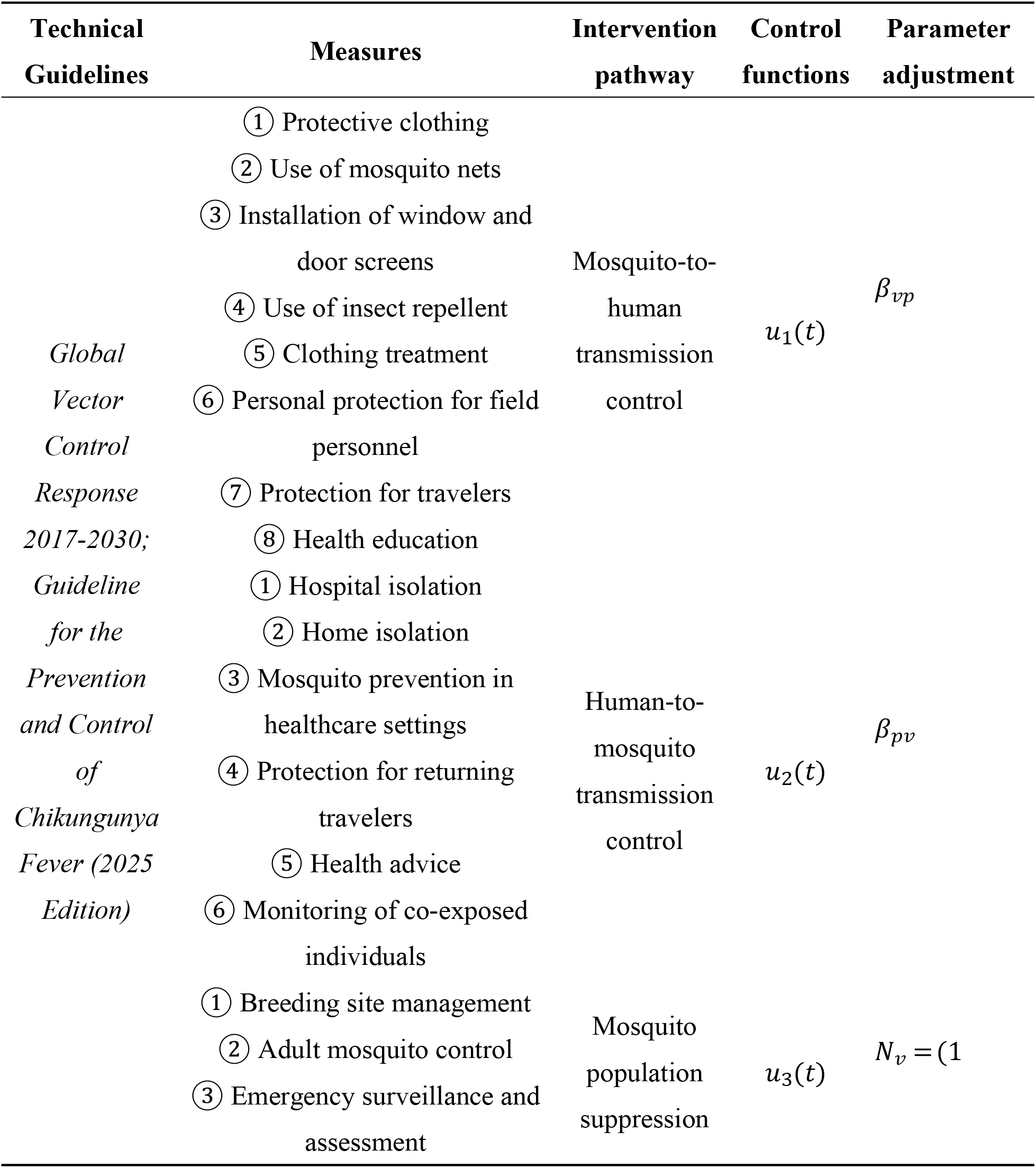
Parameterization of Intervention Measures Based on Technical Guidelines.

**Table 4.** Design of Intervention Strategy Combinations Based on Transmission Pathway Analysis.

| Strategy | Mosquito-to-human transmission control $u_1$ | Human-to-mosquito transmission control $u_2$ | Mosquito population suppression $u_3$ |
| --- | --- | --- | --- |
| Strategy 1<sup>0</sup> | 0 | 0 | 0 |
| Strategy 2 | HMC-Estimated | 0 | 0 |
| Strategy 3 | 0 | HMC-Estimated | 0 |
| Strategy 4 | 0 | 0 | HMC-Estimated |
| Strategy 5 | HMC-Estimated | HMC-Estimated | 0 |
| Strategy 6 | HMC-Estimated | 0 | HMC-Estimated |
| Strategy 7 | 0 | HMC-Estimated | HMC-Estimated |
| Strategy 8<sup>1</sup> | HMC-Estimated | HMC-Estimated | HMC-Estimated |

**Note:** <sup>0</sup> represents the blank control group with no intervention applied to any transmission pathway; <sup>1</sup> represents the control group where interventions are applied to all transmission
pathways at the actually observed intensity.

The structure of system by incorporating the above control measures was given below:

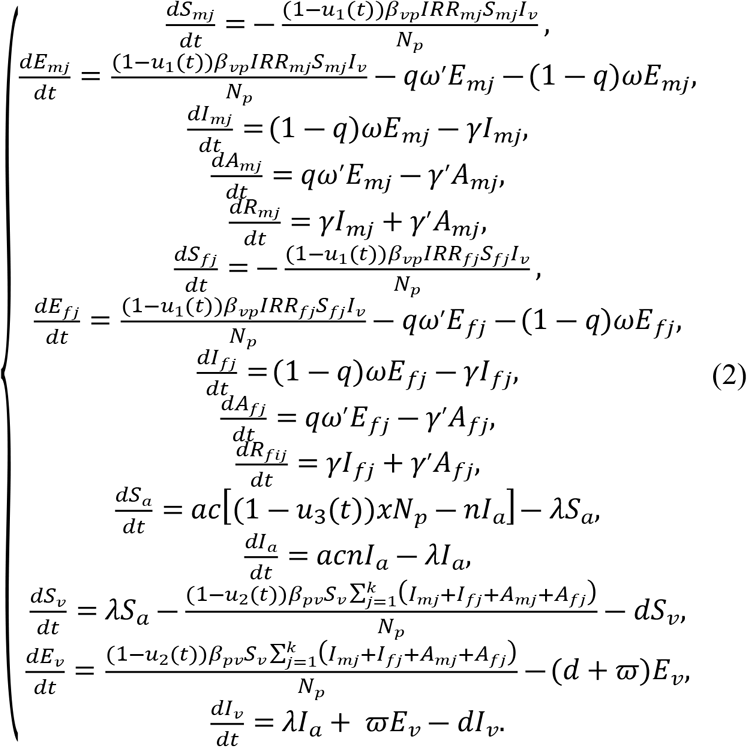

Our main objective was to minimize the number of new cases in symptomatic classes and the costs required to control epidemic. Consequently, the objective functional was defined as

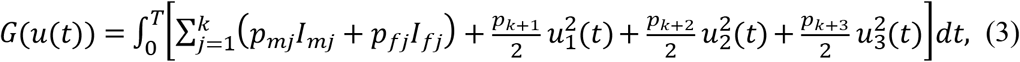

where, the constants *p*_*mj*_,*p*_*fj*_,(*j* = 1,…,*k*) are the weight factors for the symptomatic classes *I*_*mj*_,*I*_*fj*_,(*j* = 1,…,*k*), respectively, while *p*_*k*+1_, *p*_*k*+2_, and *p*_*k*+3_ are the corresponding cost factors. The core task was to determine the optimal controls for 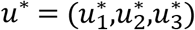 based on the Pontryagin maximum principle, such that

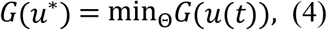

where, Θ = {*u*(*t*) ∈ *L*^3^(0,*T*)|*u*_*min*_ ≤ *u*_1_(*t*),*u*_2_(*t*),*u*_3_(*t*) ≤ *u*_*max*_, *t* ∈(0,*T*)} is the control set, T is represented the final step size, and *L*^3^(0,*T*) is the set of integrable functions defined on the interval (0,*T*).

To begin solving the optimal problem, we focused on the Lagrangian and Hamiltonian for Eqs. (2) to (4). The Lagrangian for the optimal problem is defined as follows:

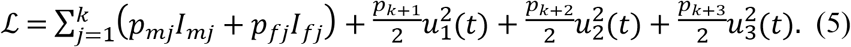

To find the minimum value of Eq. (5), we introduced the Hamiltonian function ℋ defined as follows:

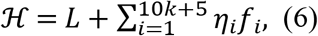

where, *f*_*i*_ is the right-hand side of the differential Eq. (2) of the i-th state variable, and *η*_*i*_ is the adjoint-vector. Based on the existence of an optimal solution for the control problem, the following Theorem 1 can be obtained.

#### Theorem 1

Let 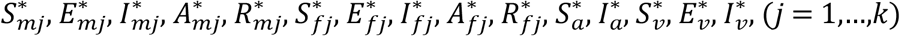 represent the state solutions associated with the optimal control measures 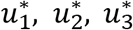 for the optimum control system stated in (2) and (3). Then, we derive the adjoint variables *η*_*i*_ (*i* = 1,…,10*k* + 5) that satisfy:

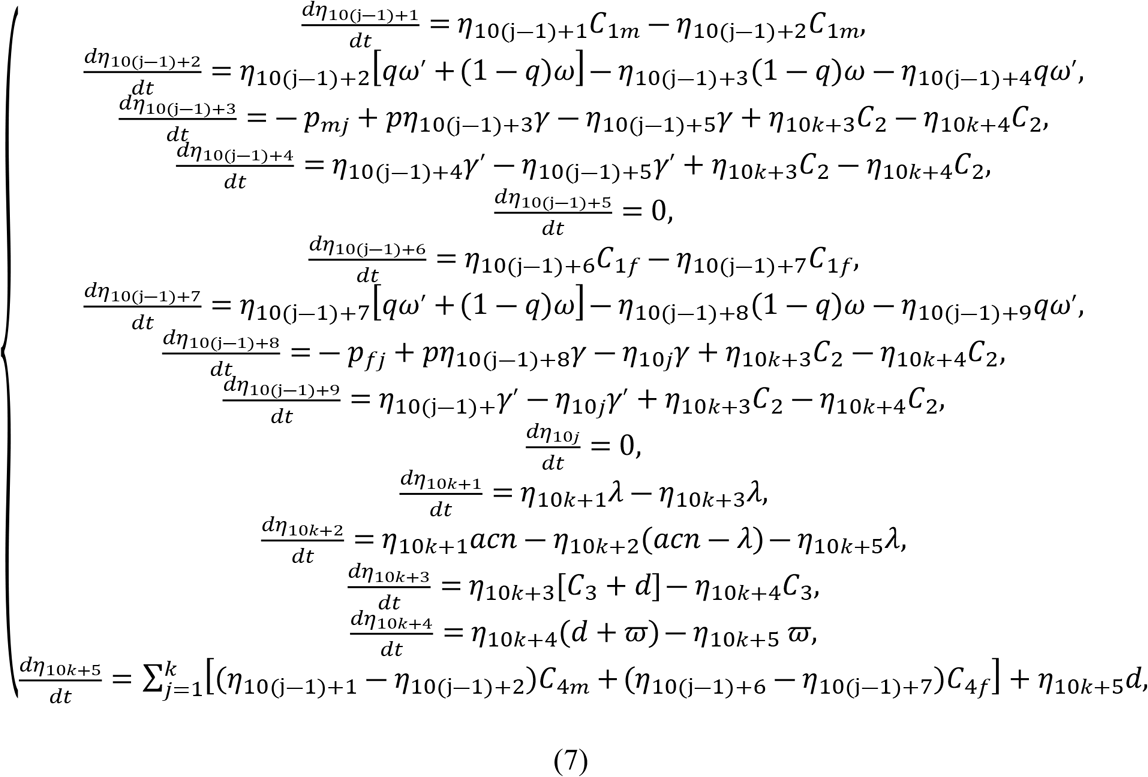

with boundary conditions or transversality conditions

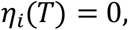

where,

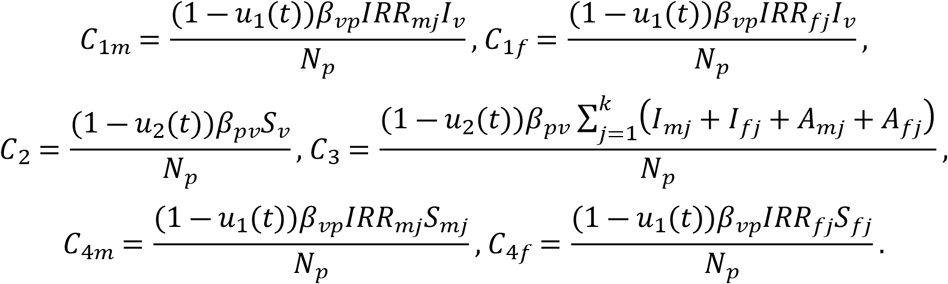

Moreover, the control measures 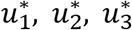 are given by

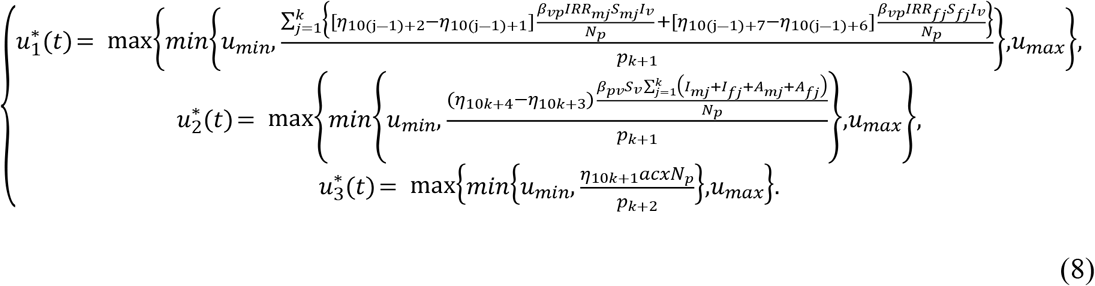

**Proof** Based on the Pontryagin maximum principle and the Hamiltonian function (Eq.(6)), the adjoint equation is obtained as follows:

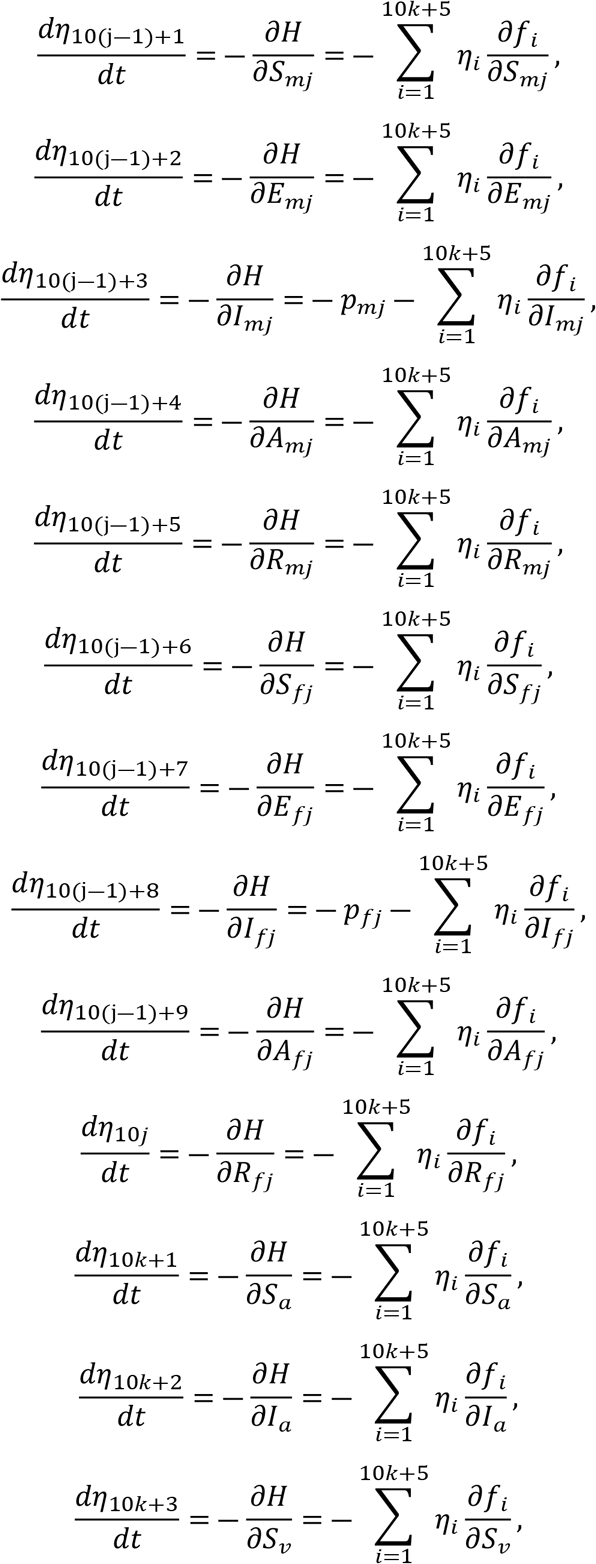

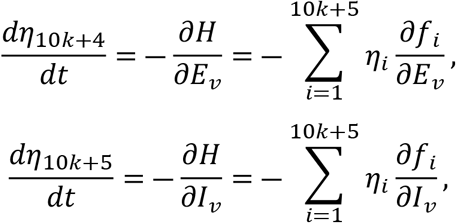

subject to boundary time conditions (i.e. final) *η*_*i*_(*T*) = 0, *i* = 1,…,10*k* + 5. In order to achieve the desired problem (8), we utilized the following equations:

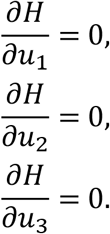

By utilizing the property of the control space Θ being in the interior of the control set, we obtained the desired result.

The theoretical framework was applied to analyze the largest recorded CHIKF outbreak in China. Considering that human interventions cannot completely interrupt transmission, the control parameters were bounded between *u*_min_ = 0.2 and *u*_max_ = 0.8. First, Model (1) was used to fit infection data from the preparedness phase, stratified into six population groups by sex and age (male: 0-14, 15-59, 60+; female: 0-14, 15-59, 60+). Model parameters were estimated using the PMCMC method (S2 Table). The estimated parameters were then applied to Model (2) for fitting the containment phase and numerically solving the control functions. Goodness-of-fit comparisons indicated that segmenting the containment phase by the incubation lag significantly improved model performance (S1 and S3 Fig).

Based on the fitting results of actual epidemic data and the control intensity curves derived from Eqs. (8) (S1 Fig), the analysis reveals that mosquito population suppression was maintained at the highest intensity (*u*_3_ = 0.8) starting from July 19. However, it was also the first measure to begin decreasing in intensity, gradually declining from September 8 and remaining at the minimum control level (*u*_3_ = 0.2) after September 11, with an average intensity of *u*_3_ = 0.7086. In comparison, both mosquito-to-human transmission control and human-to-mosquito transmission control reached their peak control intensities (*u*_1_ = *u*_2_ = 0.8) on July 20. Human-to-mosquito transmission control began to gradually decrease from September 13, dropping to its minimum (*u*_2_ = 0.2) by September 15, with an average intensity of *u*_2_ = 0.7444. In contrast, mosquito-to-human transmission control started to decline from September 15 and reached its lowest influence level (*u*_1_ = 0.2) during September 17, with an average intensity of *u*_1_ = 0.7086.

### Evaluation of Combination Strategies

In this study, we systematically evaluated the effectiveness of three intervention measures—mosquito-to-human transmission control, human-to-mosquito transmission control, and mosquito population suppression—under different implementation strategies. Using “no intervention” and “full implementation of all three measures” as reference scenarios, we simulated changes in infection numbers under single or pairwise combinations of interventions (Fig 6). Across all six population groups, the effectiveness of the strategies, ranked from highest to lowest, was as follows: “mosquito-to-human transmission control + human-to-mosquito transmission control + mosquito population suppression” (Strategy 8) > “mosquito-to-human transmission control + mosquito population suppression” (Strategy 6)> “human-to-mosquito transmission control + mosquito population suppression” (Strategy 7) > “mosquito population suppression” (Strategy 4) > “mosquito-to-human transmission control + human-to-mosquito transmission control” (Strategy 5) > “human-to-mosquito transmission control” (Strategy 3) > “mosquito-to-human transmission control” (Strategy 2) > “no control” (Strategy 1). Under the no-intervention scenario, the final epidemic size was the largest, whereas the combined implementation of all three measures resulted in the smallest final size, with an average reduction of 95.7586% in infections compared to the no-intervention scenario. The greatest reduction was observed among males aged 15-59 years (95.7620%), while the smallest reduction was seen in females aged 0-14 years (95.7518%).

**Fig 6.**
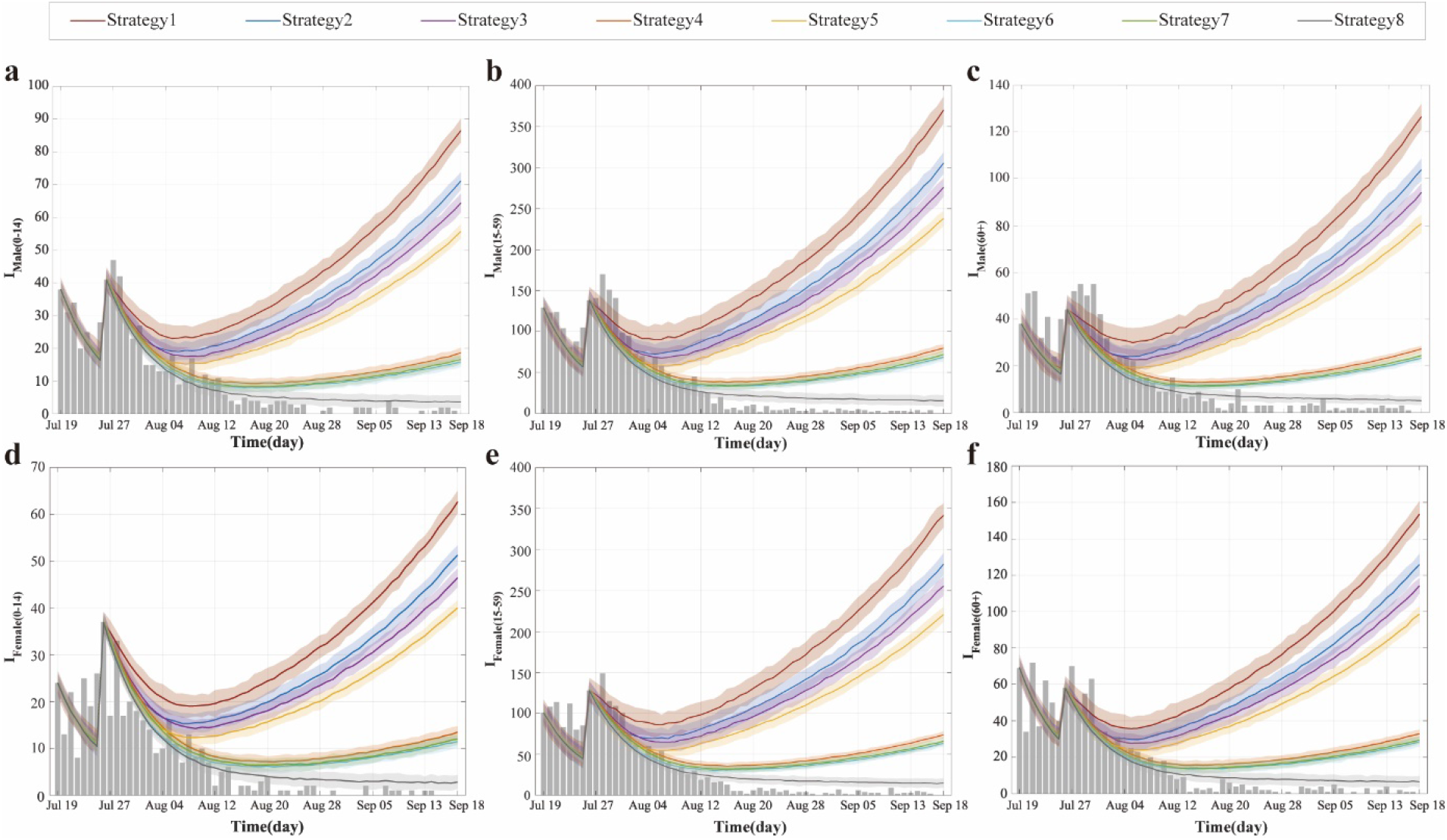
Simulated infection counts by population group under different control strategies in the largest recorded CHIKF outbreak in China. **(a-f)** Temporal dynamics of infections for six population groups: males 0-14 years (a), males 15-59 years (b), males 60+ years (c), females 0-14 years (d), females 15-59 years (e), and females 60+ years (f). Dark gray bars represent observed infections. The dark red curve (Strategy 1) represents the no-intervention control group based on segmented fitting of Model (1), while the dark gray curve (Strategy 8) corresponds to the actual control group based on segmented fitting of Model (2).

Among the single-intervention strategies, mosquito population suppression was the most effective, reducing the final epidemic size by an average of 78.4695% (range: 78.4611%-78.4764%) compared to the no-intervention scenario. This was followed by human-to-mosquito transmission control, which led to an average reduction of 25.6044% (25.5899%-25.6102%). Mosquito-to-human transmission control alone had the lowest impact, with an average reduction of 17.8302% (17.8170%-17.8365%). In terms of demographic variations, mosquito population suppression and human-to-mosquito transmission control were most effective for males aged 15-59 years, whereas mosquito-to-human transmission control performed best for females aged 0-14 years. All single measures showed the weakest effects for females aged 60 years and above. Among the two-measure combinations, mosquito-to-human transmission control + mosquito population suppression was the most effective, reducing the final epidemic size by an average of 81.5226% (81.5135%-81.5285%). This was followed by human-to-mosquito transmission control + mosquito population suppression, with an average reduction of 80.7287% (80.7208%-80.7352%). The combination of mosquito-to-human transmission control + human-to-mosquito transmission control had the lowest impact, with an average reduction of 35.7960% (35.7772%-35.7772%), which was even lower than that of mosquito population suppression alone. In terms of demographic responses, mosquito-to-human transmission control + mosquito population suppression and human-to-mosquito transmission control + mosquito population suppression were most effective for males aged 15-59 years, while mosquito-to-human transmission control + human-to-mosquito transmission control worked best for females aged 0-14 years. All two-measure combinations continued to show the weakest effects for females aged 60 years and above.

### Dose-Response Effects of Interventions

We conducted parameter sweeps for three control measures—mosquito-to-human transmission control (*u*_1_), human-to-mosquito transmission control (*u*_2_), and mosquito population suppression (*u*_3_)—by varying each within the range (0, 1) and systematically evaluated the impact of 1,331 control combinations on the effective reproduction number (ℛ_*eff*_). Results showed that across all scenarios, the ℛ_*eff*_ for the “human-to-mosquito” transmission pathway (ℛ_*eff*(*pv*)_) was consistently and significantly higher than that for the “mosquito-to-human” pathway (ℛ_*eff*(*vp*)_).

Theoretically, setting any single control parameter to 1 (i.e., implementing extreme-intensity isolation or vector control) would reduce the overall system ℛ_*eff*_ to zero, achieving complete transmission interruption. However, such extreme intensity is often impractical in real-world interventions. To identify feasible control strategies, we fixed one parameter at its actual value based on empirical data and adjusted the other two, identifying multiple non-extreme control combinations capable of reducing ℛ_*eff*_ below 1 (Fig 7a-c):

- With *u*_1_ fixed at its actual value, settings of (*u*_2_=0.6, *u*_3_=0.9), (*u*_2_= 0.8, *u*_3_=0.8), or (*u*_2_= 0.9, *u*_3_=0.6) all achieved ℛ_*eff*_=0.9737.
- With *u*_2_ fixed at its actual value, combinations (*u*_1_= 0.7, *u*_3_=0.9) or (*u*_1_= 0.9, *u*_3_ =0.7) reduced ℛ_*eff*_=0.8865.
- With *u*_3_ fixed at its actual value, combinations (*u*_1_= 0.7, *u*_2_=0.9) or (*u*_1_= 0.9, *u*_2_ =0.7) resulted in ℛ_*eff*_=0.9465.

**Fig 7.**
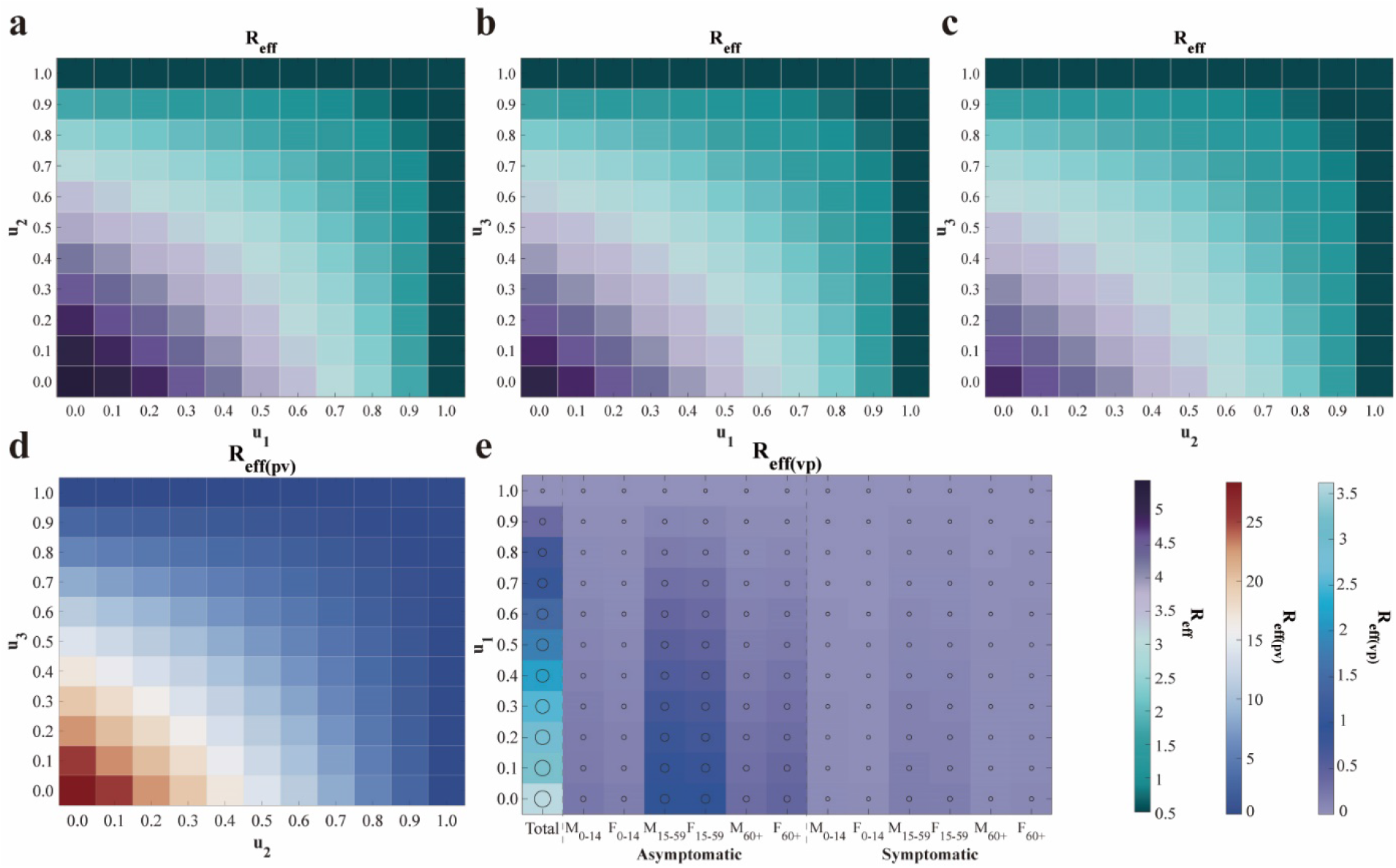
Heatmap analysis of the impact of control strategy combinations on the effective reproduction number (ℛ_*eff*_). **(a)-(c)** Effects on the overall ℛ_*eff*_ when altering combinations of the other two control measures while fixing one strategy at its actual implementation value. **(d)** Variation of the human-to-mosquito effective reproduction number (ℛ_*eff*(*pv*)_) with changes in *u*_2_ and *u*_3_. **(e)** Variation of the mosquito-to-human effective reproduction number (ℛ_*eff*(*vp*)_) and its subcomponents with changes in *u*_1_. The axes correspond to the intensities of controlling mosquito-to-human transmission (*u*_1_), human-to-mosquito transmission (*u*_2_), and mosquito population suppression (*u*_3_). Each row of subplots employs a different value range and colormap standard.

All the above outcomes satisfy ℛ_*eff*_< 1, indicating that the outbreak can be effectively controlled. Further analysis revealed that beyond these threshold combinations, increasing the intensity of any control measure further reduced ℛ_*eff*_ and enhanced control effectiveness.

Based on the mathematical structure of the basic reproduction number (ℛ_0_) and simulation results, mosquito-to-human transmission control (*u*_1_) had a relatively limited impact on the effective reproduction number for human-to-mosquito transmission (ℛ_*eff*(*pv*)_) (Fig 7d). Specifically, when *u*_2_=0.7 and *u*_3_=0.9, or *u*_2_=0.9 and *u*_3_=0.7, ℛ_*eff*(*pv*)_ remained constant at 0.8494 regardless of the value of *u*_1_. Systematically increasing the intensity of any control measure beyond these combinations further reduced the ℛ_*eff*(*pv*)_ value. On the other hand, human-to-mosquito transmission control (*u*_2_) and mosquito population suppression (*u*_3_) exhibited limited influence on the effective reproduction number for mosquito-to-human transmission (ℛ_*eff*(*vp*)_) (Fig 7e). When u_1_ reached the actual implementation level, ℛ_*eff*(*vp*)_ could be stably controlled below 1. Further simulations demonstrated that when either *u*_2_ or *u*_3_ was fixed at its actual value, increasing *u*_1_ to 0.8 was sufficient to reduce ℛ_*eff*(*vp*)_ to 0.7239, even with fluctuations in the other parameter. This indicates that transmission within the human population was effectively contained under such conditions.

Under scenarios where the intensities of all three control measures were adjusted simultaneously, the study identified multiple asymmetric strategy combinations capable of reducing ℛ_*eff*_ below the epidemic threshold. Specifically, when the intensity of one measure was maintained at 0.1 while the other two were set to 0.9, the system ℛ_*eff*_ decreased to 0.9604. Similarly, when one measure was increased to 0.9 while the other two remained at 0.7, the same control effect was achieved. More notably, when the three control intensities were specifically allocated as 0.6, 0.8, and 0.9 (excluding other numerical combinations), the system ℛ_*eff*_ was further reduced to 0.9055, indicating significant synergistic and complementary effects among the different interventions. It is important to highlight that, beyond achieving the control effects described above, further increasing the intensity of any single measure continued to lower the ℛ_*eff*_ value, demonstrating that each intervention retains potential for continuously enhancing outbreak control effectiveness.

## Discussion

This study developed a structural vector-borne model. The model system encompasses two main populations: the human population, structured using an age- and sex-stratified SEIAR compartmental framework, and the mosquito population, which includes both larval (aquatic) and adult stages. The total number of compartments in the system reaches 10*k* + 5, where *k* represents the number of age groups. Through theoretical analysis of this high-dimensional system of ordinary differential equations, the basic reproduction number (ℛ_0_) was derived. Its mathematical structure reveals that the transmission contributions from infectious individuals within the same species combine additively to form ℛ_0_, whereas the contributions from cross-species transmission (e.g., between humans and mosquitoes) are represented by their geometric mean. This finding elucidates the coupling mechanism governing interspecies transmission from a dynamical perspective.

Existing studies indicate significant age-specific disparities in the global infection burden of CHIKF. One modeling estimate suggests approximately 14.4 million annual infections worldwide, with a relatively higher burden among individuals aged 40-60 years, while extreme age groups (e.g., under 10 and over 80 years) exhibit elevated mortality risks (*5*). Furthermore, a systematic analysis of European outbreaks also identified the 45-64 age group as high-risk, with female cases generally outnumbering males (*12*). However, our analysis of the largest CHIKF outbreak in China reveals a distinct pattern: cases were slightly more prevalent among males than females, and the 15-59 age group constituted the most affected population. This discrepancy may be attributed to the relatively broad age categorization used in the current study, which led to the 15-59 group—comprising a large share of the working-age population—dominating the demographic composition. Additionally, individuals in this age range tend to have wider daily mobility and greater exposure to mosquito habitats, further elevating their infection risk. Moreover, this study found that asymptomatic infections within this group contributed substantially to transmission, suggesting their potential role as key drivers of the outbreak, which may partly explain why this demographic emerged as the core affected population in the local epidemic.

In the study of intervention strategies for vector-borne diseases, although specialized mathematical models for CHIKF remain relatively limited, research on diseases with similar transmission mechanisms, such as dengue fever, provides an important reference. Existing studies on vector-borne disease modeling have indicated that rapid containment can be achieved when surveillance, research, community engagement, and governance operate as an integrated and adaptive system (*45*). Further analysis suggests that vector control is a critical component for achieving rapid outbreak containment (*46*), and integrated approaches combining case management, community protection, and mosquito elimination have been demonstrated as the optimal strategy for interrupting virus transmission (*47-48*). The optimal control model developed in this study, based on the largest CHIKF outbreak in China data, validates these perspectives: the combined implementation of all three control measures reduced the total number of infections by an average of 95.7586%, demonstrating the significant advantage of synergistic interventions. Mosquito population suppression, as a core intervention, reached and maintained the highest intensity earliest during the containment phase and, as a single measure, led to an average reduction in infections of 78.4695%.

However, relevant studies have pointed out that unless sustained at very high implementation intensity, the effectiveness of vector control measures may gradually diminish over time, potentially leading to a resurgence of the outbreak during later phases. Premature reduction in control intensity may undermine the sustainability of intervention outcomes (*49*).

Regarding the synergistic implementation of multiple measures, this study found that the dual strategy of “mosquito-to-human transmission control + mosquito population suppression” was the most effective (reducing infection size by 81.5226%), outperforming other two-measure combinations and single interventions. This aligns with findings from dengue fever research indicating that “the combined use of mosquito nets and insecticides yields the highest infection avoidance rate” (*49*). It is noteworthy that although human-to-mosquito transmission control as a standalone measure had limited effectiveness (25.6044% reduction), its combination with mosquito population suppression significantly enhanced overall containment efficacy (80.7287% reduction). This suggests that multi-pathway transmission blocking is key to improving intervention quality when resources permit. Furthermore, the effects of different measures varied across population groups: mosquito population suppression and human-to-mosquito transmission control were most effective for males aged 15-59 years, while mosquito-to-human transmission control provided better protection for females aged 0-14 years, indicating that intervention strategies should be tailored to population-specific behavioral and exposure characteristics. The findings support the integration of surveillance-response systems, vector control, and community engagement into an adaptive framework to facilitate rapid outbreak containment.

In the phase-based analysis of epidemic control, this study divided the containment period into incubation lag and post-incubation lag subphases, revealing distinct stage-specific differences in transmission dynamics. The analysis indicated that the overall effective reproduction number (ℛ_eff_) during the incubation lag subphase was slightly lower than that in the subsequent subphase, with both “mosquito-to-human” transmission and population-specific contributions remaining relatively low in the earlier subphase. It is noteworthy that asymptomatic individuals aged 15-59 years consistently served as the core transmission group across both subphases, underscoring their pivotal role in outbreak control. These findings align with transmission models that incorporate time-delay effects (*50*), suggesting that the transmission delay induced by the incubation period should be fully considered in intervention assessments. The study further revealed that during the post-incubation lag subphase, the overall system ℛ_eff_ was 1.3243—still above the epidemic threshold—yet the “mosquito-to-human” transmission intensity (0.8122) had fallen below 1, indicating a potential gradual decline in human transmission. However, the “human-to-mosquito” transmission intensity remained elevated at 2.1593, reflecting persistent viral activity in mosquito populations and a continued risk of outbreak resurgence.

To effectively contain the spread of the outbreak, this study systematically evaluated the effectiveness of different control strategies, demonstrating that multiple non-extreme control combinations could reduce ℛ_eff_ below the epidemic threshold. When fixing one intervention at its actual implementation level while optimizing the other two, effective strategies consistently achieved ℛ_eff_<1 across scenarios: with u_1_ fixed, combinations such as (*u*_2_=60%, *u*_3_=90%), (*u*_2_=80%, *u*_3_=80%), or (*u*_2_=90%, *u*_3_=60%) yielded ℛ_eff_=0.9737; with u_2_ fixed, combinations like (u_1_=70%, u_3_=90%) reduced ℛ_eff_ to 0.8865; and with *u*_3_ fixed, configurations such as (*u*_1_=70%, *u*_2_=90%) lowered ℛ_eff_ to 0.9465. Further analysis revealed distinct pathway-specific effects: mosquito-to-human transmission control (*u*_1_) exhibited limited influence on human-to-mosquito transmission (ℛ_eff(pv)_), which remained stable at 0.8494 when *u*_2_ and *u*_3_ were set to 70% and 90% respectively, whereas mosquito-to-human transmission (ℛ_eff(vp)_) was primarily controlled by *u*_1_, reaching 0.7239 when its intensity attained 80% even under variations in the other measures. When simultaneously adjusting all three interventions, specific asymmetric intensity allocations—such as single-measure intensity at 10% paired with dual-measure intensity at 90%, or a triple-measure profile of 60%/80%/90%—achieved ℛ_eff_ values as low as 0.9055. All control measures demonstrated continuous effectiveness enhancement, where increasing intensity beyond baseline levels consistently yielded further reduction in transmission risk.

This study has several limitations. First, with the recent approval and rollout of the world’s first CHIKF vaccine, Ixchiq, by the U.S. FDA in 2023 and its subsequent authorization in multiple countries (*51*), vaccination has become an important intervention strategy. However, the model developed in this study only considers non-pharmaceutical interventions and does not incorporate the role of immunization strategies within the overall control framework. Second, the current model does not fully account for the potential impact of climate change on mosquito distribution and virus transmission. Studies have shown that global warming and altered precipitation patterns are significantly expanding the suitable habitats for *Aedes* mosquitoes (*52-54*), increasing transmission risks in temperate regions (*55*). Yet, the present model does not integrate meteorological variables to reflect this dynamic process. Furthermore, the model inadequately addresses the role of human mobility. In the context of globalization, international travel and cross-regional movement accelerate virus spread (*56-57*). However, the model is primarily based on local transmission dynamics and does not systematically analyze the impact of imported cases on local outbreaks. Future research should aim to integrate vaccination strategies, climate variables, and human mobility data, combined with genomic approaches, to establish a more comprehensive risk assessment framework.

## Conclusions

The optimal control analytical framework developed in this study effectively enables the analysis of mosquito-borne outbreak transmission. Asymptomatic individuals among males aged 15-59 years were identified as the core drivers of transmission in the largest CHIKF outbreak recorded in China in 2025. Mosquito population suppression should be prioritized as a universal core intervention. In light of population heterogeneity, it is recommended to strengthen the interruption of human-to-mosquito transmission and mosquito density control for males aged 15-59 years, while focusing on mosquito-to-human transmission protection for females aged 0-14 years. For females aged 60 years and above, multi-pathway comprehensive intervention strategies should be adopted to compensate for the limited effectiveness of single measures in this demographic. These findings provide a scientific basis for developing population- and phase-specific precision control strategies, and hold practical significance for enhancing emergency response capacity for mosquito-borne diseases.

## Author contributions

Conceptualization: T. M. C., J. H. L., Z. Y. Z., J. R. Investigation: J. H. L., Z. Y. Z., J. R., J. G. Z., Q. X. L, W. T. S., Q. P. C. Methodology: J. H. L., Z. Y. Z., J. R., S. P., R. F. Software: J. H. L., J. G. Z., Q. X. L. Validation: Z. Y. Z., J. R., K. G. L., S. P., R. F., Y. H. S., T. X. X. Writing—original draft: J. H. L., J. G. Z., Q. X. L., K. G. L. Writing—review & editing: J. H. L., Z. Y. Z., J. R., S. P., R. F., Y. H. S., Q. P. C., T. M. C., T. X. X. All authors read and approved the final manuscript.

